# Supergene evolution triggered by the introgression of a chromosomal inversion

**DOI:** 10.1101/234559

**Authors:** Paul Jay, Annabel Whibley, Lise Frezal, Angeles de Cara, Reuben W. Nowell, Jim Mallet, Kanchon K. Dasmahapatra, Mathieu Joron

## Abstract

Supergenes are groups of tightly linked loci whose variation is inherited as a single Mendelian locus and are a common genetic architecture for complex traits under balancing selection^1^. Supergene alleles are long-range haplotypes with numerous mutations underlying distinct adaptive strategies, often maintained in linkage disequilibrium through the suppression of recombination by chromosomal rearrangements^2–5^. However, the mechanism governing the formation of supergenes is not well understood, and poses the paradox of establishing divergent functional haplotypes in face of recombination^1,6^. Here, we show that the formation of the supergene alleles encoding mimicry polymorphism in the butterfly *Heliconius numata* is associated with the introgression of a divergent, inverted chromosomal segment. Haplotype divergence and linkage disequilibrium indicate that supergene alleles, each allowing precise wing-pattern resemblance to distinct butterfly models, originate from over a million years of independent chromosomal evolution in separate lineages. These “superalleles” have evolved from a chromosomal inversion captured by introgression and maintained in balanced polymorphism, triggering supergene inheritance. This mode of evolution is likely to be a common feature of complex structural polymorphisms associated with the coexistence of distinct adaptive syndromes, and shows that the reticulation of genealogies may have a powerful influence on the evolution of genetic architectures in nature.

---

How new beneficial traits which require more than one novel mutation emerge in natural populations is a long-standing question in biology^7–9^. Supergenes control alternative adaptive strategies that require the association of multiple co-adapted characters, and have evolved repeatedly in many taxa under balancing selection. Examples include floral heteromorphy determining alternative pollination strategies^10^, butterfly mimicry of alternative wing-pattern and behaviours of toxic models^3,11,12^, contrasting mating tactics in several birds^13,14^, and alternative social organization in ant colonies^4^. In most documented cases, the maintenance of character associations is mediated by polymorphic rearrangements, such as inversions, which suppress local recombination and allow the differentiated supergene alleles to persist^2,4,5,10,13^. However, the build-up of differentiated haplotypes from initially recombining loci is poorly understood^1,6^. Recombination is necessary to bring into linkage mutations that arise on different haplotypes, but also acts to break down co-adapted combinations. While inversions may capture epistatic alleles at adjacent loci, this requires adaptive polymorphism at both loci prior to the rearrangement. Third, linkage disequilibrium around functional mutations under balancing selection persists only over short evolutionary times^15^. The few models of supergene evolution^7,16^ do not readily yield the conditions for the formation of differentiated haplotypes or the evolutionary trajectory of functional genetic elements within rearranged non-recombining regions after the initial structural variation.

To understand allelic evolution in supergenes, we studied Amazonian populations of the butterfly *Heliconius numata,* in which up to seven distinct wing-pattern morphs coexist (Fig. 1a), each one matching to near perfection the colours and shapes of other toxic Lepidoptera (Heliconiinae, Danainae, Pericopiinae)^2^. This balanced polymorphism is controlled by a supergene locus (*P*) associated with an inversion polymorphism^2^ that captures multiple genetic loci controlling wing-pattern variation in butterflies and moths^12,18–21^ and allows multiple wing elements to be inherited as a single Mendelian character. The ancestral chromosomal arrangement, called *Hn0*, is associated with the recessive supergene allele^17^ which controls the widely distributed morph *silvana*. All other characterized supergene alleles, grouped into a family of alleles called *Hn1*, determine a diversity of mimetic morphs dominant to *silvana* and associated with the 400-kb inversion P_1_ (Fig. 1a; Ref. ^2,17^). A subset of these alleles is associated with additional rearrangements (P_2_) in adjacent positions^2^. The emergence of the P supergene architecture is therefore associated with the evolution of inversion P_1_ polymorphism, followed by adjacent rearrangements. P_1_ forms a well-differentiated haplotype distinct from the ancestral haplotype along its entire length (Fig. 4a and Supplementary Fig. 3), and with extreme values of linkage disequilibrium (LD)^2^. This inversion therefore stands as a block of up to 7000 differentiated SNPs along its 400 kb length, associated with supergene evolution, adaptive diversification and dominance variation.

**Figure 1.**
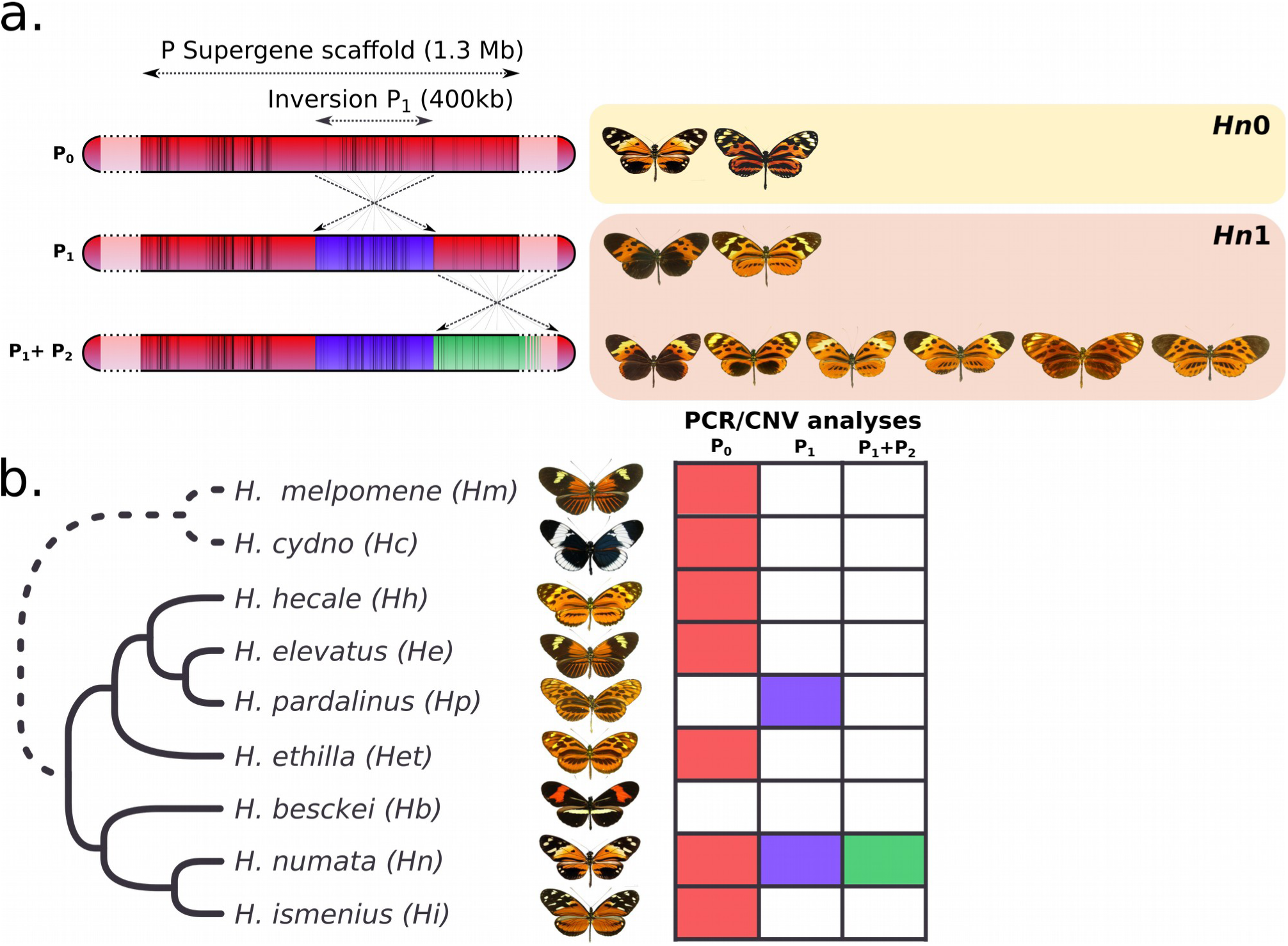
Distribution of supergene inversions in the silvaniform clade of *Heliconius*. **a** Structure of the *H. numata* (*Hn*) mimicry supergene *P* characterised by polymorphic inversions and some of the morphs associated with each arrangement. *P* allows *Hn* to produce highly distinguishable morphs in the same location. The first derived inversion (P_1_, blue), is common to all rearranged alleles (*Hn1*), and distinguishes them from the ancestral, recessive *P* alleles (mimetic forms *silvana* or *laura*, *Hn0*). The P dominant allele (Andean mimetic form *bicoloratus* and *peeblesi*) is controlled by a rearrangement including only the chromosomal inversion P_1_. A further rearrangement (P_2_, green) linked to the first inversion is associated with a large diversity of derived, intermediate dominant mimicry alleles^2,17^. A 4kb duplication was also detected only in individuals showing the inversion P_1_. **b** Presence/absence of the two major rearrangements in species closely-related to *H. numata* (silvaniform clade), tested by PCR of breakpoint-diagnostic markers, and independently by duplication-diagnostic CNV assays. All species are fixed for the ancestral arrangement (red), except *H. pardalinus* (*Hp*), fixed for P_1_, and *H. numata* showing polymorphism for P_1_ and P_2_.

*Heliconius numata* belongs to the so-called silvaniform clade of ten species which diverged ca. 4 My ago from its sister clade (Fig. 1b; Fig. 2a; Supplementary Fig. 1; Ref. ^22^). To explore the origin and evolution of inversion P_1_, we investigated the presence of this inversion in other species of the clade. PCR amplification of inversion breakpoints showed that inversion *P*_1_ was polymorphic in *H. numata* (*Hn*) across its Amazonian range, and was also found fixed in all population of *H. pardalinus* (*Hp*), a non-sister species deeply divergent from *H. numata* within the silvaniform clade (Fig. 1b and Supplementary Table 3). All other taxa including the sister species of *H. numata* and that of *H. pardalinus* were positive only for markers diagnostic of the ancestral gene order. Furthermore, a 4kb duplication associated with P_1_ in *Hn* was also found in whole genome sequence datasets for all *Hp* individuals and no other taxon (Fig. 1b; Supplementary Table 2; Supplementary Fig. 10). Breakpoint homology and similar molecular signatures in *Hp* and *Hn* are thus consistent with a single origin of this inversion. This sharing of P_1_ between non-sister species could be due to the differential fixation of an ancient polymorphic inversion (incomplete lineage sorting, ILS), or to a secondary transfer through introgression.

**Figure 2.**
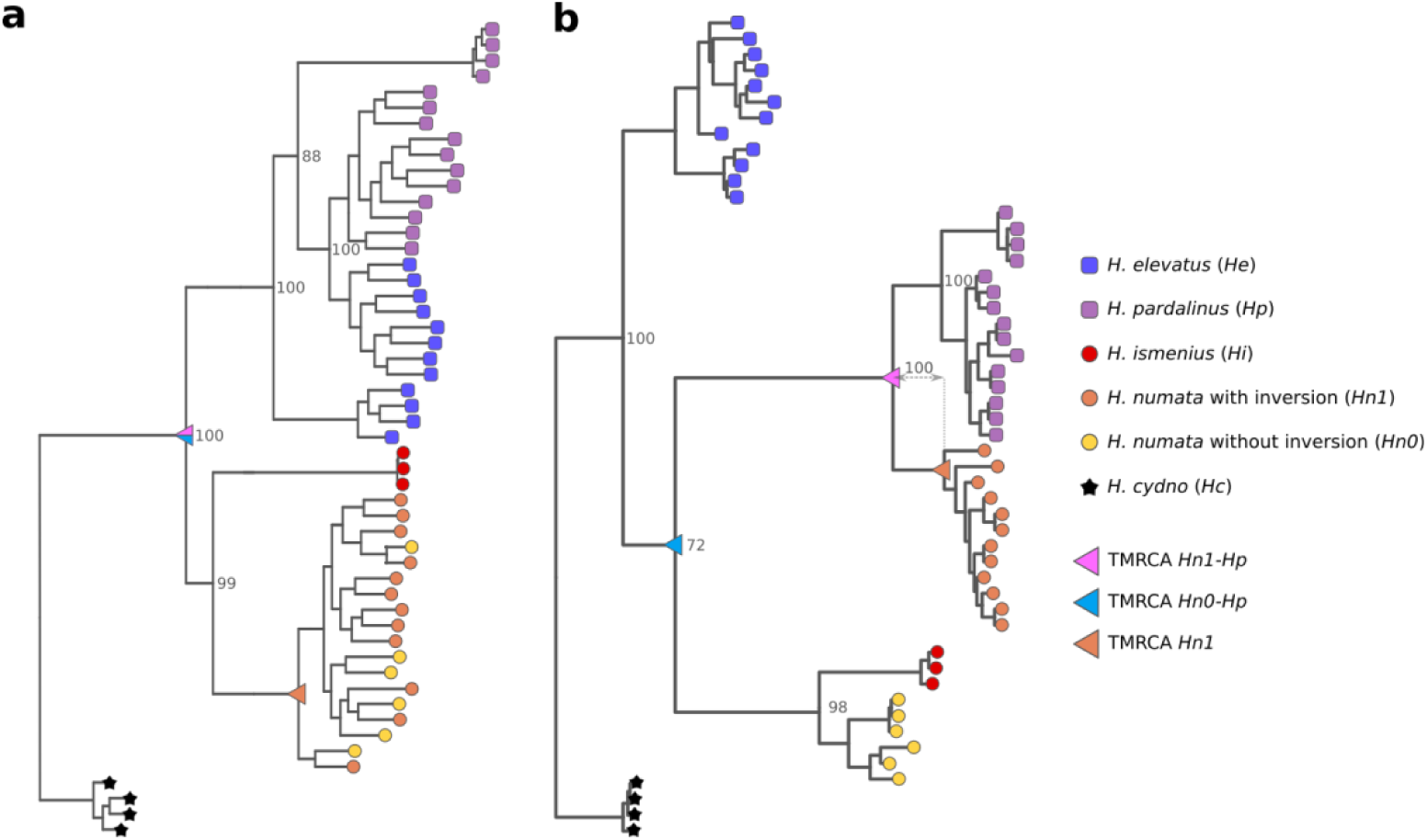
Whole genome and inversion phylogenies of *H. numata* and related species. **a** Whole genome phylogeny, showing two well-separated branches grouping *H. pardalinus* and *H. elevatus* on the one hand and *H. numata* and *H. ismenius* on the other hand, consistently with previous studies (*i.e*. Ref. ^22^. See Supplementary Fig. 1 for the phylogeny with all taxa). **b** Undated inversion P_1_ phylogeny. All *Hn* individuals displaying the inversion P_1_ (*Hn1*) group with *Hp*, while *Hn* individuals displaying the ancestral arrangement (*Hn0*) remain with sister species *Hi. He* groups closer to the outgroup (*Hc*) reflecting introgression with *H. melpomene*, a species closely related to *Hc* (Supplementary Fig. 1; Ref. ^23^). For clarity, only species informative to introgression history are represented here; The inversion is a 400kb segment displaying much phylogenetic heterogeneity among the other taxa, reflecting a complex history of gene flow and incomplete lineage sorting (see in Supplementary Fig. 1 for phylogenies including all taxa).

To clarify whether this sharing between *Hp* and *Hn* is a rare anomaly, specifically associated with the supergene locus, or is a common feature that is also found elsewhere in the genome, we estimated the excess of shared derived mutations between sympatric *Hp* and *Hn*, relative to an allopatric control, *H. ismenius* (*Hi*, sister species of *Hn*), using the *fd* statistic^24^. We estimated that a significant 6.2 % of the genome was shared via gene flow between *Hn* and *Hp* (mean *fd* = 0.062*. Fig. 3 and Supplementary Fig. 2c), consistent with a general signal of genome-wide gene flow between *Hn* and other species within the silvaniform clade (Supplementary Fig. 2) and between other *Heliconius* species^23^. When *fd* is estimated using *Hn* specimens homozygous for inversion P_1_ (*Hn1*), the supergene scaffold is associated with a strong peak of shared derived mutations between *Hn* and *Hp* (Fig. 3, blue arrow). This is not observed between *Hn1* and other silvaniforms (Supplementary Fig. 2) nor when using *Hn* specimens homozygous for the ancestral supergene arrangement (*Hn0*; Supplementary Fig. 2c). Between *Hn1* and *Hp*, the entire P_1_ inversion shows a high level of *fd*, which drops to background levels precisely at inversion breakpoints (Fig. 4b). *Hn1* and *Hp* therefore share a block of derived mutations associated with the inversion.

**Figure 3.**
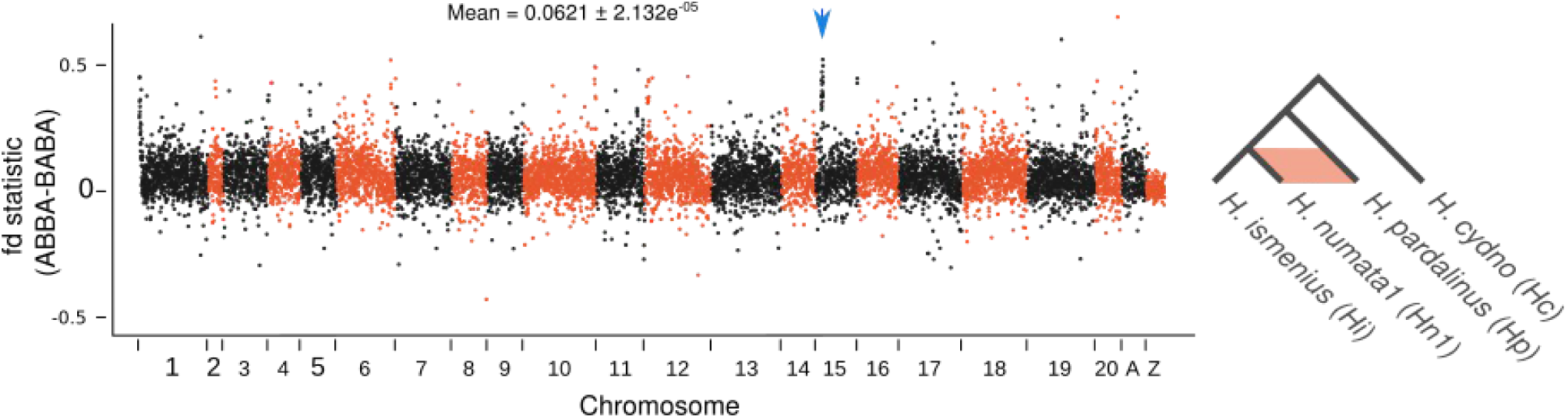
Excess of shared derived mutations between *Hp* and Hn1. *fd* statistic computed in non-overlapping 20 kb sliding window and plotted along the whole genome. The ABBA-BABA framework and related statistics assess the excess of shared derived mutations between *Hp* and *Hn1*, relative to a control (*Hi*) not connected by gene flow to the others. Outgroup *Hc* allows the mutations to be polarized. A mean *fd* = 0 is expected if *Hp* is not connected by gene flow to *Hn1*. Unmapped contigs are grouped within an “A” chromosome. The supergene scaffold (HE667780) is indicated by a blue arrow. Standard error was assessed with block jackknifing (600 kb block size).

**Figure 4.**
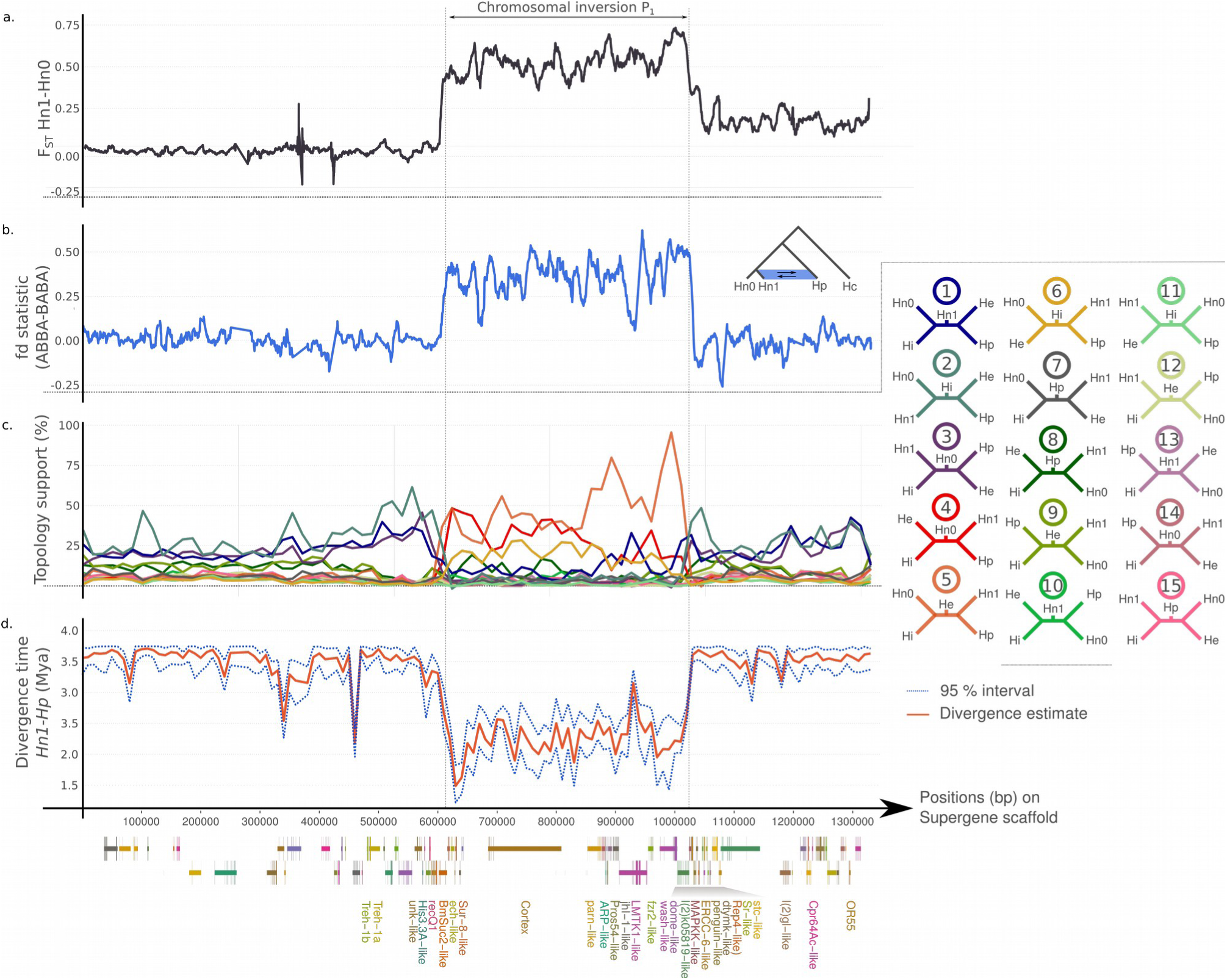
Phylogenetic and divergence variation at the supergene scaffold. **a** F_ST_ scan between Hn1 and Hn0. Inversion P_1_ shows a generally high Fst value contrary to the rest of the genome. P_2_ rearrangement (1028-1330 kb) shows lower but nonetheless elevated F_st_ values. **b** *fd* statistic (ABBA-BABA) computed in 10 kb sliding windows (increment = 500bp) with P1= *Hn0*, *P2*=*Hn1*, *P3*=*Hp*, *O*=*Hc*. Outside the inversion, a *fd* value close to 0 is observed, as expected under a no gene flow scenario. At P_1_ inversion breakpoints, the *fd* values strongly increase and remains high in the whole inversion. **c** Weightings (Twisst^25^) for all fifteen possible phylogenetical topologies between *H. numata* with inversion (*Hn1*), *H. numata* without inversion (*Hn0*), *H. ismenius* (*Hi*), *H. pardalinus* (*Hp*) and *H. elevatus* (*He*), with loess smoothing (level = 0.05).Topology 2 is the species topology. Strong topology change occurs around inversion breakpoints. Whithin the inversion, the best supported topologies (4, 5 & 6) group *Hn1* close to *Hp*. **d** Variation in divergence time between *Hn1* and *Hp*, computed in 10 kb non-overlapping sliding windows. The divergence time inside the inversion is significantly lower than in the rest of the genome.

Because high values of *fd* in a given genome region may be due to gene flow or to incomplete lineage sorting, we estimated the divergence times of *Hn1* and *Hp* within and outside of inversion P_1_. The unique ancestor of inversion P_1_ in *Hp* and *Hn1* was estimated to be 2.30 My old (95% interval 1.98-2.63 Mya, Fig. 4d; Pink triangles in Fig. 2), significantly more recent than the divergence time of the rest of the genome (3,59 Mya; 95% interval 3.37-3.75 Mya) (Fig. 4d and Supplementary Fig. 4), which indicates that the inversion was shared among lineages well after their split. Introgression may be dated to an interval between the time to the most recent common ancestor (TMRCA) of *Hp* and *Hn1* inversions (*i.e*. 2.30 Mya, Fig 4d; Pink triangles in Fig. 2) and the TMRCA of all *Hn1* inversions (2.24 Mya, 95 % interval 1.89-2.59 Mya, Supplementary Fig. 5d; Orange triangle in Fig. 2), i.e. about 1.30 My after *Hp-Hn* speciation. To estimate the age of the inversion, we identified the two sequences of the 4kb duplicated region associated with the inversion in an *Hn1* BAC library and in an *Hn1* genome assembly, and estimated their divergence time. We found that the duplication occurred 2.41 Mya (95% interval 1.96-2.71 Mya), indicating that P_1_ may have spread between lineage *Hp* and *Hn* shortly after the occurrence of the inversion.

To determine the direction of introgression, we surveyed the position of the sister species to *Hn* (*Hi*) and to *Hp* (*H. elevatus*, *He*) in phylogenies computed along the supergene scaffold and in other regions of the genome. The genome as a whole and regions flanking the inversion all show a similar topology to the one found by Kozak *et al*. ^22^, with expected sister relationships of *Hi* and *Hn* and of *He* and *Hp* (Fig. 2a). Evaluating the support for each possible topology among the five informative taxa (*Hn0*, *Hn1*, *Hi*, *Hp*, *He*) using Twisst^25^ confirmed the consistent support for the separation of (*Hn*, *Hi*) and (*Hp*, *He*) clades despite a high level of incomplete lineage sorting within each clade (Fig. 4c and Supplementary Figs 6-7). By contrast, the inversion P_1_ shows strong support for topologies that group *Hn1* with *Hp*, and major topology changes coincide with inversion breakpoints (Figs 2b and 4c and Supplementary Figs 6-8), consistent with a single origin of the inversion. Within the inversion, the highest support consistently goes to *Hn1* grouping within (*Hp*, *He*) and away from (*Hn*, *Hi*) (Fig. 4c, topology 5), indicating an introgression from *Hp* to *Hn*. This conclusion is robust to the species used as sister groups to *Hn* or *Hp* (Supplementary Fig. 6d-g). Alternative topologies (4 and 6) are also found in relatively high proportions in the interval ~650-850kb, presumably owing to high levels of incomplete lineage sorting at the clade level in this region, or ancient gene flow among other species of the clade. Supporting these interpretations, topology analysis with taxa unaffected by *Hn1-Hp* introgression (for instance using *Hn0* and replacing *Hp* with a closely related species, *H. hecale*) still showed the same pattern of unresolved phylogenetic signal in this interval between the two major branches of the clade (*Hp-He-H*. *hecale* vs. *Hn-Hi*) (Supplementary Fig. 6h-i) suggesting that the lack of support to a given topology is independent of and predates the introgression. This segment is centred on the gene *cortex*, known to control wing pattern variation in several lepidopteran species and showing a rich history of selective sweeps and introgression among Heliconius taxa^19,20,26^. The topological variation observed at the supergene may therefore reflect a combination of incomplete lineage sorting and gene flow at different time periods. This suggests that segmental inversions, like speciation, can capture and preserve phylogenetic variations inherited from the entire history of gene flow and selection among recombining taxa.

Sustained differentiation between *P* alleles over the entire length of the inversion in *H. numata* is therefore explained by the 1.3 My of independent evolution of an inverted haplotype within *H. pardalinus*, whose divergence was maintained and accentuated after introgression by the suppression of recombination. Our results show that, as previously hypothesised^1,6,13^, complex balanced polymorphism such as those controlled by supergene may evolve via the differentiation of rearranged haplotypes in separate lineages, followed by adaptive introgression in a host population where differentiated haplotypes are preserved through suppression of recombination, and maintained by balancing selection. This provides the first empirical evidence for a mechanism to explain the formation of supergene, and offers a parsimonious solution to the paradox of the evolution of divergent haplotypes in face of recombination.

Supergene formation through adaptive introgression requires an initial selective advantage to the inversion in the recipient population, and balancing selection maintaining the polymorphism. In *H. numata*, the introgressed arrangement is associated with a successful melanic phenotype (*bicoloratus*) mimicking abundant local species in the foothills of the Andes and enjoying a 7-fold increased protection relative to ancestral arrangements^27^. This introgression likely constitutes an ecological and altitudinal expansion to premontane Andean foothills where the melanic wing mimicry ring dominates, and an empirical example for the theoretical role of inversions as “adaptive cassettes” triggering eco-geographical expansions in an introgressed lineage^28^. Despite their role in reproductive isolation^29^, inversions may be prone to adaptive introgression through combined selection on linked mutations^30^. This is supported by the rapid introgression of inversion P_1_ after it was formed.

Inversion P_1_ linked with the adjacent rearrangement P_2_, is also associated with other well-protected mimetic forms^2,27^, and most *H. numata* phenotypes associated with the inversion are unmatched in *H. pardalinus*, indicating that introgression was followed by further adaptive diversification to local mimicry niches. Balancing selection, mediated by negative assortative mating among inversion genotypes, prevents the fixation of the inversion, as reflected by a deficit of homozygotes for the introgressed haplotype in the wild^31^. Supergene evolution is therefore consistent with the introgressed inversion having a strong advantage under mimicry selection but being maintained in a polymorphism with ancestral haplotypes by negative frequency-dependence.

These findings demonstrate that introgression, especially when involving structural variants, may trigger the emergence of novel genetic architectures. This scenario may underlie the evolution of many complex polymorphisms under balancing selection in a wide variety of organisms, such as the MHC loci in vertebrates^32^, self-incompatibility systems in plants^33^, mating types in fungi^34^ or, much more generally, sex chromosomes. Our results therefore shed new light on the importance of introgression as a mechanism shaping the architecture of genomes and assisting the evolution of complex adaptive strategies.

## Methods

### Sampling and sequencing

Specimens of *H. numata*, *H. ismenius*, *H. elevatus*, *H. pardalinus*, *H. hecale*, *H. ethilla*, *H. besckei*, *H. melpomene* and *H. cydno* were collected in the wild in Peru, Ecuador, Colombia, French Guiana, Panama and Mexico (Supplementary Table 1). Bodies were conserved in NaCl saturated DMSO solution at −20°C and DNA was extracted using Qiagen DNeasy blood and tissue kits according to the manufacturers’ instructions and with RNase treatment. Illumina Truseq paired-end whole genome libraries were prepared and 2×100bp reads were sequenced on the Illumina HiSeq 2000 platform. Reads were mapped to the *H. melpomene* Hmel1 reference genome^23^ using Stampy v1.0.23^35^ with default settings except for setting the substitution rate to 0.05 to allow for expected divergence from the reference. Alignment file manipulations used SAMtools v0.1.19^36^. After mapping, duplicate reads were excluded using the *MarkDuplicates* tool in Picard (v1.107; http://broadinstitute.github.io/picard) and local indel realignment using IndelRealigner was performed with GATK v2.1.5^37^. Invariant and polymorphic sites were called with GATK UnifiedGenotyper. Filtering was performed on individual samples using GATK VariantFiltration to remove sites with depth <10 or greater than 4 times the median coverage of the sample, or sites with low mapping quality (using the expression “MQ < 40.0 || MQ0>= 4 && ((MQ0 /(1.0*DP))>0.1)”. SnpSift filter^38^ was used to exclude sites with QUAL or GQ less than or equal to 30. After filtering, variant call files were merged using GATK CombineVariants.

### PCR analysis and genotyping

Inversion breakpoints were genotyped by PCR amplification of genomic DNA using Thermo Scientific^®^ Phusion High-Fidelity DNA Polymerase. Primer sequences and PCR conditions used are specified in Supplementary Table 4.

### Duplication Analysis

Copy number analysis of the supergene scaffold was performed on resequence alignments after duplicate removal and local realignment using CNVnator v0.3^39^ with default settings and a bin size of 100bp.

The 4kb sequence detected as duplicated was blasted^40^ against the Hn1 BAC clone library from Ref. 2 and against a *H. numata* genome, generated by the Heliconius consortium using a combination of SMRT long read (Pacific Biosciences) and Illumina short read (Discovar assembly), and available on LebBase (http://ensembl.lepbase.org). Three BAC clones (38g4, 24i10 and 30F8) and two scaffolds (scaffold13474 and scaffold16807) showed high blast values (e-value=0). Their entire sequences were mapped on the *H. melpomene* reference genome^23^ with BLAST. They correspond to two regions close to the two breakpoints of inversion P_1_. The sequences resulting from the duplications were extracted from the BAC clones and the scaffolds and aligned with MUSCLE^41^.

### ABBA-BABA analysis

ABBA-BABA analyses were conducted with the scripts provided by Ref. ^42^. The *fd* statistic was computed in 20 kb non-overlapping windows for the whole genome (min. genotyped position=1000) and 10 kb sliding windows with a 500bp step, (min. genotyped position=500) for the supergene scaffold (HE667780). Block Jackknife (600 kb block) resampling was performed to assess the significance of whole genome *fd* values.

### Phylogenetic analyses

To determine the direction of introgression, we use the fact that the introgressed species should appear phylogenetically closer than expected to the donor species, but also closer to the sister species of the donor. Thus, considering a species topology like (A,B),(C,(D,E)), a sequence showing a (A,C)(D,(E,B)) topology probably arose by the way of an introgression from E to B, whereas a sequence showing a ((B,E),A)(D,C) topology probably arose via introgression from B to E. To search for such patterns, we computed a whole genome phylogeny and several phylogenies at different locations within and outside the inversion.

The whole genome phylogeny was obtained with SNPhylo^43^, with 100 bootstraps and *H. cydno* as the outgroup. RAxML^44^was used to determine local phylogenies, with GTRCAT model and 100 bootstrap. Nevertheless, we found that individuals from the different species were frequently mixed and the species topology was highly variable, complicating the interpretation of topology changes at the inversion location. We thus used Twisst^25^ to unravel the changes in topology and assess phylogenetic discordance along the supergene scaffold. We used Beagle^45^ to phase the haplotypes of the supergene scaffold, with 10000 bp size and 1000 bp overlapping sliding windows. Maximum likelihood trees were generated with the phyml_sliding_window.py script with the GTR model and a 50 SNP sliding window (https://github.com/simonhmartin/twisst).

### Divergence time analyses

To discriminate between introgression and ancestral polymorphism hypotheses, Bayesian inferences of the divergence time between *H. pardalinus* and *H. numata* were made with Phylobayes^46^ Analyses were performed on 10 kb non-overlapping sliding windows, using all individuals of the two species and including individuals of all other species in our dataset to obtain better resolution.Date estimates were relative to the divergence of *H. cydno* with the silvaniform clade, estimated by Ref. ^22^ to be approximately 3,84 Mya., using a log-normal autocorrelated relaxed clock. Each chain ran for at least 30000 states, with 10000 burn-in states. Chain convergence was checked with Tracer (http://beast.bio.ed.ac.uk/Tracer). Resultant trees and time estimates were analysed with ete3 python library^47^.

Divergence of the duplication-associated sequences was done in the same way. Whole genome resequence data from all specie except *Hn1* and *Hp* were used, as well as sequences from the three BAC clones and the *H. numata* genome. *Hn1* and *Hp* specimens were not used, as they tend to artificially increase the mutation rate inferred by Phylobayes (cause to the duplication, *Hn1* and *Hp* appear highly heterozygous at the duplication location. See supplementary Fig. 10)

## References

1. Schwander, T., Libbrecht, R. & Keller, L. Supergenes and Complex Phenotypes. Curr. Biol. 24, R288–R294 (2014).

2. Joron, M. et al. Chromosomal rearrangements maintain a polymorphic supergene controlling butterfly mimicry. Nature 477, 203–206 (2011).

3. Kunte, K. et al. doublesex is a mimicry supergene. Nature 507, 229–232 (2014).

4. Wang, J. et al. A Y-like social chromosome causes alternative colony organization in fire ants. Nature 493, 664–668 (2013).

5. Lamichhaney, S. et al. Structural genomic changes underlie alternative reproductive strategies in the ruff (Philomachus pugnax). Nat. Genet. 48, 84–88 (2016).

6. Llaurens, V., Whibley, A. & Joron, M. Genetic architecture and balancing selection: the life and death of differentiated variants. Mol. Ecol. 26, 2430–2448 (2017).

7. Charlesworth, D. & Charlesworth, B. Theoretical genetics of Batesian mimicry II. Evolution of supergenes. J. Theor. Biol. 55, 305–324 (1975).

8. Fisher, R. A. The Genetical Theory Of Natural Selection. (At The Clarendon Press, 1930).

9. Franks, D. W. & Sherratt, T. N. The evolution of multicomponent mimicry. J. Theor. Biol. 244, 631–639 (2007).

10. Li, J. et al. Genetic architecture and evolution of the S locus supergene in Primula vulgaris. Nat. Plants 2, 16188 (2016).

11. Timmermans, M. J. T. N. et al. Comparative genomics of the mimicry switch in Papilio dardanus. Proc R Soc B 281, 20140465 (2014).

12. Joron, M. et al. A Conserved Supergene Locus Controls Colour Pattern Diversity in Heliconius Butterflies. PLOS Biol. 4, e303 (2006).

13. Tuttle, E. M. et al. Divergence and Functional Degradation of a Sex Chromosome-like Supergene. Curr. Biol. 26, 344–350 (2016).

14. Küpper, C. et al. A supergene determines highly divergent male reproductive morphs in the ruff. Nat. Genet. 48, 79–83 (2016).

15. Charlesworth, D. Balancing selection and its effects on sequences in nearby genome regions. PLoS Genet. 2, e64 (2006).

16. Yeaman, S. Genomic rearrangements and the evolution of clusters of locally adaptive loci. Proc. Natl. Acad. Sci. 110, E1743–E1751 (2013).

17. Le Poul, Y. et al. Evolution of dominance mechanisms at a butterfly mimicry supergene. Nat. Commun. 5, 5644 (2014).

18. Huber, B. et al. Conservatism and novelty in the genetic architecture of adaptation in Heliconius butterflies. Heredity 114, 515–524 (2015).

19. Nadeau, N. J. et al. The gene cortex controls mimicry and crypsis in butterflies and moths. Nature 534, 106–110 (2016).

20. van’t Hof, A. E. et al. The industrial melanism mutation in British peppered moths is a transposable element. Nature 534, 102–105 (2016).

21. Van Belleghem, S. M. et al. Complex modular architecture around a simple toolkit of wing pattern genes. Nat. Ecol. Evol. 1, 0052 (2017).

22. Kozak, K. M. et al. Multilocus Species Trees Show the Recent Adaptive Radiation of the Mimetic Heliconius Butterflies. Syst. Biol. 64, 505–524 (2015).

23. The Heliconius Genome Consortium. Butterfly genome reveals promiscuous exchange of mimicry adaptations among species. Nature 487, 94–98 (2012).

24. Martin, S. H., Davey, J. W. & Jiggins, C. D. Evaluating the Use of ABBA–BABA Statistics to Locate Introgressed Loci. Mol. Biol. Evol. 32, 244–257 (2015).

25. Martin, S. H. & Van Belleghem, S. M. Exploring evolutionary relationships across the genome using topology weighting. Genetics 206, 429–438 (2017).

26. Enciso-Romero, J. et al. Evolution of novel mimicry rings facilitated by adaptive introgression in tropical butterflies. Mol. Ecol. 26, 5160–5172 (2017).

27. Chouteau, M., Arias, M. & Joron, M. Warning signals are under positive frequency-dependent selection in nature. Proc. Natl. Acad. Sci. U. S. A 113, 2164–2169 (2016).

28. Kirkpatrick, M. & Barrett, B. Chromosome inversions, adaptive cassettes and the evolution of species’ ranges. Mol. Ecol. 24, 2046–2055 (2015).

29. Hoffmann, A. A. & Rieseberg, L. H. Revisiting the Impact of Inversions in Evolution: From Population Genetic Markers to Drivers of Adaptive Shifts and Speciation? Annu. Rev. Ecol. Evol. Syst. 39, 21–42 (2008).

30. Kirkpatrick, M. & Barton, N. Chromosome inversions, local adaptation and speciation. Genetics 173, 419–434 (2006).

31. Chouteau, M., Llaurens, V., Piron-Prunier, F. & Joron, M. Polymorphism at a mimicry supergene maintained by opposing frequency-dependent selection pressures. Proc. Natl. Acad. Sci. 114, 8325–8329 (2017).

32. Grossen, C., Keller, L., Biebach, I., International Goat Genome Consortium & Croll, D. Introgression from domestic goat generated variation at the major histocompatibility complex of Alpine ibex. PLoS Genet. 10, e1004438 (2014).

33. Castric, V., Bechsgaard, J., Schierup, M. H. & Vekemans, X. Repeated adaptive introgression at a gene under multiallelic balancing selection. PLoS Genet. 4, e1000168 (2008).

34. Corcoran, P. et al. Introgression maintains the genetic integrity of the mating-type determining chromosome of the fungus Neurospora tetrasperma. Genome Res. 26, 486–498 (2016).

35. Lunter, G. & Goodson, M. Stampy: a statistical algorithm for sensitive and fast mapping of Illumina sequence reads. Genome Res. 21, 936–939 (2011).

36. Li, H. et al. The Sequence Alignment/Map format and SAMtools. Bioinforma. Oxf. Engl. 25, 2078–2079 (2009).

37. DePristo, M. A. et al. A framework for variation discovery and genotyping using next-generation DNA sequencing data. Nat. Genet. 43, 491–498 (2011).

38. Cingolani, P. et al. Using Drosophila melanogaster as a Model for Genotoxic Chemical Mutational Studies with a New Program, SnpSift. Front. Genet. 3, 35 (2012).

39. Abyzov, A., Urban, A. E., Snyder, M. & Gerstein, M. CNVnator: An approach to discover, genotype, and characterize typical and atypical CNVs from family and population genome sequencing. Genome Res. 21, 974–984 (2011).

40. Altschul, S. F., Gish, W., Miller, W., Myers, E. W. & Lipman, D. J. Basic local alignment search tool. J. Mol. Biol. 215, 403–410 (1990).

41. Edgar, R. C. MUSCLE: multiple sequence alignment with high accuracy and high throughput. Nucleic Acids Res. 32, 1792–1797 (2004).

42. Martin, S. H., Davey, J. W. & Jiggins, C. D. Evaluating the use of ABBA-BABA statistics to locate introgressed loci. Mol. Biol. Evol. msu269 (2014).

43. Lee, T.-H., Guo, H., Wang, X., Kim, C. & Paterson, A. H. SNPhylo: a pipeline to construct a phylogenetic tree from huge SNP data. BMC Genomics 15, 162 (2014).

44. Stamatakis, A. RAxML version 8: a tool for phylogenetic analysis and post-analysis of large phylogenies. Bioinformatics 30, 1312–1313 (2014).

45. Browning, S. R. & Browning, B. L. Rapid and Accurate Haplotype Phasing and Missing-Data Inference for Whole-Genome Association Studies By Use of Localized Haplotype Clustering. Am. J. Hum. Genet. 81, 1084–1097 (2007).

46. Lartillot, N., Lepage, T. & Blanquart, S. PhyloBayes 3: a Bayesian software package for phylogenetic reconstruction and molecular dating. Bioinformatics 25, 2286–2288 (2009).

47. Huerta-Cepas, J., Serra, F. & Bork, P. ETE 3: Reconstruction, Analysis, and Visualization of Phylogenomic Data. Mol. Biol. Evol. 33, 1635–1638 (2016).

